# Genetic Influences on Hormonal Markers of Chronic HPA Function in Human Hair

**DOI:** 10.1101/055244

**Authors:** Elliot M. Tucker-Drob, Andrew Grotzinger, Daniel A. Briley, Laura E. Engelhardt, Frank D. Mann, Megan Patterson, Clemens Kirschbaum, Emma K Adam, Jessica A. Church, Jennifer Tackett, K.Paige Harden

## Abstract

Cortisol is the primary output of the hypothalamic-pituitary-adrenal (HPA) axis and is central to the human biological stress response, with wide-ranging effects on physiological function and psychiatric health. In both humans and animals, cortisol is frequently studied as a biomarker for exposure to environmental stress. Relatively little attention has been paid to the possible role of genetic variation in heterogeneity in chronic cortisol, in spite of well-studied biological pathways of glucocorticoid function. Using recently developed technology, hair samples can now be used to measure accumulation of cortisol over several months. In contrast to more conventional salivary measures, hair cortisol is not influenced by diurnal variation or transient hormonal reactivity. In an ethnically and socioeconomically diverse sample of 1 070 child and adolescent twins and multiples from 556 unique families, we estimated genetic and environmental influences on hair concentrations of cortisol and its inactive metabolite, cortisone. We identified sizable genetic influences on cortisol that decrease with age, concomitant with genetic influences on cortisone that increase with age. Shared environmental influences on cortisol and cortisone were modest and, for cortisol, decreased with age. Twin-specific, non-shared environmental contributions to cortisol and cortisone became increasingly correlated with age. We find some evidence for sex differences in the biometric contributions to cortisol, but no strong evidence for main or moderating effects of family socioeconomic status on cortisol or cortisone. This study constitutes the first genetic study of hormone concentrations in human hair, and provides the most definitive characterization to-date of age and socioeconomic influences on hair cortisol.

## Genetic Influences on Hormonal Markers of Chronic HPA Function in Human Hair

The biological stress system produces a multifaceted regulatory response to physiological and psychological threats to homeostasis.^1^ Within seconds of stressor onset, the hypothalamic-pituitary-adrenal (HPA) axis releases corticotropin-releasing hormone from the hypothalamus, stimulating the pituitary to release adrenocorticotropic hormone, which in turn stimulates release of glucocorticoids (specifically, cortisol, in humans) from the adrenal cortex. Glucorticoids have wide-ranging effects on physiology, including suppressing immune function and gonadal function, stimulating cardiovascular function, and elevating blood glucose.^2,3^ Moreover, glucocorticoids pass into the central nervous system, where there are several neural regions containing high densities of glucocorticoid receptors.^4^

As the key end product of the HPA-axis stress response with highly active effects on brain function, cortisol has become a leading candidate physiological mechanism for the effects of both chronic and acute stress on psychiatric health and psychopathology. HPA dysregulation, as indexed by cortisol concentrations in serum, saliva, or urine, has been linked with chronic stressors and history of major trauma,^5^ and has been concurrently and prospectively associated with a range of psychiatric symptomologies and disorders, including depression, anxiety, and posttraumatic stress disorder.^4,6,7^ Animal models that experimentally manipulate the social environment have found that HPA dysregulation is associated with brain atrophy and with suppression of signals for neurogenesis and synapse formation.^8^ Moreover, in animal models, administering exogenous glucocorticoids produces similar deleterious effects on neural structure.^4^

## Overcoming Methodological Challenges in Measuring HPA Function using Hair Sampling

That cortisol output follows a pattern of diurnal variation complicates research on the role of HPA function in chronic stress and psychopathology. Diurnal variation typically begins with cortisol levels rising in the early morning, peaking immediately after waking, and declining through the day and evening. Cortisol levels measured in blood and saliva closely track diurnal patterns of output, and urinary levels reflect output over a period of 12–48 hours.^9^ Because of this diurnal variation, single samples of cortisol in bodily fluids confound temporally stable individual differences in basal cortisol levels with within-person fluctuations. Researchers, therefore, typically attempt to index chronic levels of HPA function by taking repeated salivary samples across the day and over multiple days,^10^ a costly approach that imposes participant burden, and carries the risk of participant non-compliance and dropout. Although repeated measurements of cortisol across the day are necessary for estimating elements of the diurnal rhythm such as the cortisol awakening response or diurnal cortisol decline,^11^ hair cortisol measurement provides a more efficient and accurate assessment of long-term average or basal cortisol output.

Recently, researchers have developed methods for the analysis of cortisol accumulations in hair samples collected noninvasively at the time of the laboratory visit.^12^ Hair cortisol captures the accumulation of free cortisol over several months.^9^ Internal consistency estimates as estimated using duplicate sampling are above .90,^13^ month long test-retest consistencies are above .80,^14^ and yearlong test-retest consistencies are over .70.^15^ Hair cortisol concentrations have been estimated to correspond at over .60 with estimates of total cortisol output from thrice daily sampling of saliva taken over a one month period,^14^ but convergent validity is much lower for urinary sampling^14,16^ and for salivary estimates taken over periods of 3–4 days,^17^ which are currently considered best practices. Hair cortisol concentrations are robust to a number of possible confounds, including natural hair color, oral contraceptive use, smoking, use of everyday hair products, and frequency of hair washes.^18^ Hair cortisol is associated, as expected, with known disrupters of normal HPA functions, including shift work, Cushing syndrome, and posttraumatic stress disorder.^19^ Some studies have also reported a negative association between hair cortisol and socioeconomic status in children^20-22^ and in adults.^23^ However, null associations between socioeconomic status and hair cortisol have also been reported.^24,25^

### Unclear Role of Genetic Variation in HPA Axis Function

Although variation in HPA axis output is most commonly discussed as a biomarker for exposure to environmental stress, genetic variation is also a potential contributor to heterogeneity in HPA axis output.^4,5,26^ Indeed, the biological pathways from gene sequence to cortisol production, reception, and regulation are well-studied, and polymorphisms in these cortisol-relevant genes may account for heterogeneity in HPA function and cortisol output.^27–29^ However, biomarkers of HPA function are not currently available in samples of genotyped individuals that are sufficiently large for a well-powered genetic association study of quantitative traits.^30,31^

For many psychiatrically-relevant phenotypes, only a small subset of the specific genetic polymorphisms that constitute genetic risk have been identified, but vast literatures from twin and family designs provide precise, replicable estimates of heritability and genetic covariance with other phenotypes. For HPA axis function, however, even this basic information is lacking, as there has been relatively little research from genetic epidemiology on glucocorticoid output. A 2003 review identified only 12 genetically-informed studies of cortisol.^32^ These studies were limited by failures to account for the cortisol diurnal rhythm, small sample sizes (no study exceeded 150 twin pairs), and a lack of methodological consistency. There have been a handful of more recent twin studies of salivary cortisol, most notably a study of 700 individuals from 309 twin families,^33^ and another study of 446 twin pairs.^34^ Overall, heritability estimates of salivary cortisol have been moderate (∼30–40%), with the largest heritability estimates found for samples taken at waking. Whether these estimates, which may be downwardly biased by transient fluctuations in cortisol, generalize to overall cortisol output over a period of months is an open question.

Genetic influences on chronic HPA function may vary across subgroups. In animal models, marked sex differences exist in HPA reactivity, in the direction of greater HPA activity in females compared to males. Moreover, sex hormones modulate HPA function.^35,36^ Low socioeconomic status has also been linked to HPA dysregulation,^37^ and genetically-influenced heterogeneity in reaction norms to stressful socioeconomic contexts (i.e., a “diathesis-stress” pattern) would be expected to produce a link between low socioeconomic status and increased heritability of cortisol.^38^ Age has been reported to be associated with HPA function, in the direction of greater glucocorticoid output in adolescents compared to children and adults.^35^ Age may also modulate genetic influences on HPA activity, because of genetically-influenced variation in physiological acclimation to long-term stressors over time.^5^ Finally, genetic influence on HPA function may be activated by biological changes associated with puberty, and by stressful challenges associated with navigating social transitions across development. Overall, the extent to which individual differences in chronic HPA axis function reflect genetic differences between people, and the extent to which this genetic influences vary across subgroups, is critical information for researchers attempting to understand the relationships between genotype, chronic environmental stress, and psychiatric outcomes.

### Goals of the Current Study

Using data from an ethnically and socioeconomically diverse population-based sample of over 1 000 3^rd^ to 12^th^ grade twins, the current article reports results from a genetic epidemiological study of influences on long-term HPA function, as indexed with endocrine assays of hair.^15^ We examine both cortisol, the primary active glucocorticoid in humans, and cortisone. In humans, cortisol is metabolized into the inactive cortisone form, which can, in turn, be converted back to active cortisol to modulate glucocorticoid activity.^20,39,40^ We estimate main effects of biological sex, family socioeconomic status, and age on hair cortisol and cortisone, and moderating effects of these three factors on the genetic and environmental components of hair cortisol and cortisone variation and covariation. This is, to our knowledge, the first genetic epidemiological study of hair markers of neuroendocrine hormones, the largest twin study of neuroendocrine concentrations in any tissue type, and among the largest studies of neuroendocrine concentrations in hair to date (see^20,25,41^ for other large-scale studies of hair cortisol in singletons).

## Method

Twin pairs were recruited from the Texas Twin Project,^42^ an ongoing study of school-age twins and multiples and their parents, residing in the Austin and Houston, Texas metropolitan areas. Participants ranged in age from 7.80 to 19.47 years of age (*M* = 12.42, *SD* = 2.78). Two female participants reported endocrine disorders and were excluded from analyses. The final sample consisted of 1 141 individuals forming 607 pairs from 556 unique families. In order for a pair to be included in the current analyses, at least one member must have provided a usable hair sample. Of the 1 141 individuals in the sample, 1 070 individuals provided usable hair samples for cortisol and cortisone assay. Of the 607 pairs included, there were 533 pairs in which both members provided usable hair samples. Two families had two sets of twins, one family had quadruplets who contributed six pairwise contributions, 21 families had triplets that contributed three pairwise comparisons, and two families had triplets where only one triplet provided hair resulting in two pairwise combinations. The final sample consisted of 188 monozygotic (MZ) pairs (110 female, 78 male) and 419 dizygotic (DZ) pairs (114 female, 82 male, 223 opposite-sex). Sixty-five percent (65%) of the sample was non-Hispanic white, 5% of participants were African American, 18% of participants were Hispanic, and 12% of participants were another race/ethnicity or multiple race/ethnicities. Of the participating families, 34% reported receiving some form of means-tested public assistance, including food stamps, since the twins' birth.

### Measures

*Zygosity*. Opposite-sex pairs were classified as DZ. For same-sex pairs, zygosity was assessed using a questionnaire concerning the twin’s physical similarities (e.g., facial appearance) and the frequency that they are mistaken for one another. Twins over 14 years old completed the zygosity survey, and at least one parent and two research assistants completed the survey for all twin pairs. Responses from all raters were entered into a latent class analysis (LCA) to obtain the above classifications. LCA of physical similarity ratings has been reported to accurately determine zygosity greater than 99% of the time, as validated by genotyping.^43^

*Hair Steroid Analyses*. Hair samples were collected to determine cortisol and cortisone concentrations. On the day of the appointment, participants were instructed not to use any hair products that are not rinsed out of the hair. Samples were only collected if the participants’ hair was at least 3 cm in length. A section of hair strands approximately 3 mm in diameter was cut as close to the scalp as possible from a posterior vertex position (i.e., the center of the back of the head). Samples were analyzed at the laboratory of one of the authors (CK) using liquid chromatography-tandem mass spectrometry, as previously described.^15^ The 3 cm hair segment closest to the scalp was used for analyses. This hair segment is taken to represent cortisol and cortisone secretion over the most recent 3–month period.^15^

*Socioeconomic Status (SES)*. Years of parental education were averaged together and standardized; log-income was standardized; and the transformed education and income variables were averaged and standardized to create an SES composite.

### Analyses

To correct positive skew, log and square root transformations were applied to cortisol and cortisone, respectively. Outliers were separately winsorized for males and females by replacing extreme values with the highest observed scores within 3 *SDs* of the mean. This involved replacing 19 female and 8 male outliers for cortisol, and 10 female and 11 male outliers for cortisone. Outcomes were residualized for the year the hair samples were assayed to control for batch effects. Finally, the transformed winsorized cortisol and cortisone values were standardized (relative to the respective standard deviations of their residuals from regressions of cortisol and cortisone on age and age-squared).

Models were estimated with full information maximum likelihood using *Mplus^44^* For descriptive statistics, phenotypic models were fit using the complex sampling option to correct standard errors for nesting of individuals within families. For the biometric models, the complex sampling option was used to correct standard errors for the dependency between sibling pairs within triplet and quadruplet sets.

## RESULTS

### Descriptive Statistics

Sex differences in age trends were tested by regressing cortisol and cortisone on age, sex, and an age × sex interaction. For cortisol, there was a significant effect of age,*β* =.12, SE = .04,*p* = .001; and an age × sex interaction, *β*= −.19, SE = .06, *p* = .002. Similarly, for cortisone there was a significant effect of age, *β* =.19, SE = .06*p* < .001; and an age × sex interaction, *β*= –.20, SE = .06, *p* = .001. Females had lower levels of cortisol and cortisone relative to males at age 8, but female hormone levels increased at a faster rate over age such that concentrations were slightly higher in females than in males by age 18 (Figure 1).

**Figure 1.**
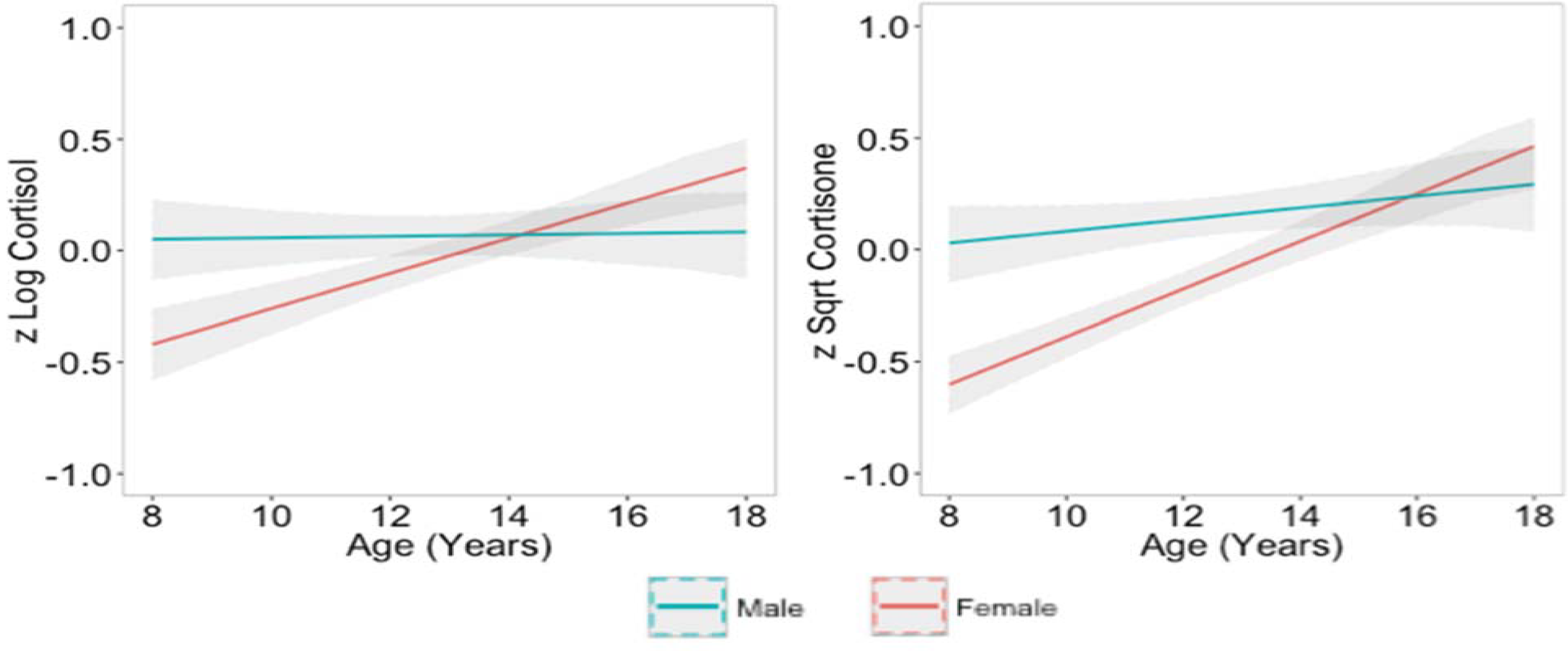
Sex-specific age trends in mean levels of cortisol and cortisone. Cortisol and cortisone were log and square root transformed, respectively, and standardized. Gray bands indicate 95% confidence intervals.

Sex-specific phenotypic correlations are summarized in Table 1. The phenotypic correlation between cortisol and cortisone concentrations was high for males (.51, 95% Confidence Interval [CI]: .43, .59) and females (.54, 95% CI: .48, .60), and largely unaffected when controlling for sex-specific age trends. The correlation between SES and both hormones was minimal for males and females. Intraclass correlations for cortisol and cortisone, split by zygosity and sex, are reported in Table 2. MZ intraclass correlations were larger than DZ intraclass correlations.

**Table 1.** Phenotypic correlations (lower diagonal) and partial correlations (upper diagonal) between cortisol, cortisone, SES, and age.

|  | Male |  |  | Female |  |  |
| --- | --- | --- | --- | --- | --- | --- |
|  | Cortisol | Cortisone | SES | Cortisol | Cortisone | SES |
| Cortisol | - | .51 (.43, .59) | .01 (-.08, .11) | - | .51 (.45, .57) | .04 (-.04, .11) |
| Cortisone | .51 (.43, .59) | - | -.07 (-.16, .03) | .54 (.48, .60) | - | .00 (-.07, .08) |
| SES | .02 (-.06, .10) | -.06 (-.15, .03) | - | .02 (-.08, .12) | -.01 (-.09, .07) | - |
| Age | .01 (-.06, .08) | .06 (-.01, .14) | - | .23 (.15, .30) | .30 (.23, .36) | - |

*Note.* Correlations on the upper diagonal controlled for sex-specific linear and quadratic effects of age. 95% confidence intervals are given in parentheses.

**Table 2.** Cross-twin intraclass correlations for cortisol and cortisone by sex and zygosity.

|  | Monozygotic |  | Dizygotic |  |  |
| --- | --- | --- | --- | --- | --- |
|  | Male | Female | Male | Female | Opposite-sex |
| Cortisol | .74 (.55, .93) | .58 (.44, .73) | .53 (.34, .72) | .34 (.17, .51) | .26 (.12, .41) |
| Cortisone | .55 (.40, .71) | .49 (.35, .63) | .47 (.33, .62) | .36 (.19, .52) | .22 (.04, .40) |

*Note.* Phenotypic correlations were estimated from models that controlled for sex-specific linear
and quadratic effects of age. 95% confidence intervals are given in parentheses.

Race/ethnicity differences in hormone levels were tested by entering three dummy-coded variables into a linear regression with Caucasian participants as the reference group. There was a significant effect of African-American race on higher cortisol (*β* =.20, SE = .06,*p* < .001), but not for Hispanic ethnicity (*β* = .03, SE = .04, *p* = .44) or for the Other/multiple race/ethnicity variable (*β* =−.01, SE = .04,*p* = .90). There was a significant effect of Hispanic ethnicity on higher cortisone (*β* =.13, SE = .05, *p* = .01), but not of African-American race (*β* =.07, SE = .05, *p* = .14) or Other/multiple race/ethnicity (*β* =.09, SE = .05, *p* = .08).

### Moderated Biometric Models

We estimated a series of three-group (MZ, same-sex DZ, opposite-sex DZ) bivariate correlated factors models (Figure S1) to estimate additive genetic (A), shared environmental (C),and non-shared environmental (E) variance components. See the Supplement for additional information on model specification.

***Moderation by sex***. We tested for *qualitative* sex differences, in which different genes influence the phenotype in males versus females, and *quantitative* sex differences, in which the same genes influence the phenotype but the magnitude of genetic influences differ. Fit indices and parameter estimates are summarized in Table S1. There was no evidence of qualitative sex differences in cortisol or cortisone, but there was evidence for quantitative sex differences in cortisone. Table S2 reports unstandardized parameter estimates from the model that included quantitative sex differences in cortisone. Across males and females, 65% of the total variability in cortisol was explained by additive genetic effects. For cortisone, additive genetic and nonshared environmental effects both explained large portions of variability for males (*h*^2^ = 44% and *e*^2^ = 35%) and females (*h*^2^ = 47% and *e*^2^ = 47%). Shared environmental effects accounted for 21% of the variance in cortisone in males, but only 6% in females.

***Moderation by age***. Model fits and model-implied parameter estimates at younger (10 years) and older (16 years) ages are reported in the top portion of Table S3. The full age moderation model was the best fitting model. Unstandardized parameter estimates for this model are reported in Table S4, and Figure 2 depicts model-predicted age trends in the *ACE* influences on cortisol, cortisone, and their bivariate association (see Figure S2 for proportions of variance).

**Figure 2.**
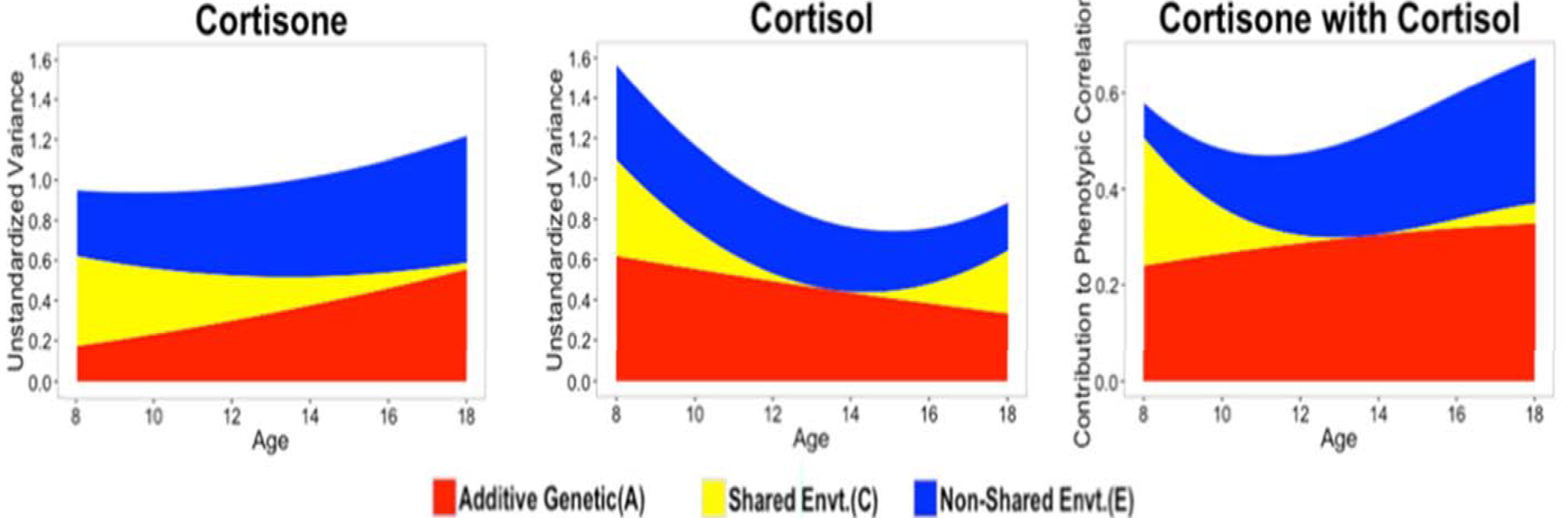
Model-implied age trends in additive genetic (A), shared environmental (C), and nonshared environmental (E) contributions to variance in cortisol, cortisone and their bivariate association.

For cortisol, non-shared environmental influences were relatively constant across the age range, whereas additive genetic effects gradually decreased with age. Shared environmental influences on cortisol decreased through age 14, at which point they fixated at zero. Although parametric results seem to indicate a re-emergence of shared environment influences in the late teenage years, nonparametric analyses indicated no such re-emergence (Figure S4). For cortisone, the additive genetic factor explained increasing variability with age, and shared environmental factor explained decreasing variability with age that were estimated near 0 by age 18. The effect of non-shared environment on cortisone was moderate and relatively stable across the age range. Across the age range, the association between cortisol and cortisone was approximately 50% attributable to shared genetic etiology. The remaining cortisol-cortisone covariation shifted from being shared environmentally mediated in the younger, preadolescent years, to being nonshared environmentally mediated in the older, adolescent years.

***Moderation by SES***. Model fits and model-implied parameter estimates at lower (1 *SD* below the mean) and higher (1 *SD* above the mean) levels of SES are reported in the top portion of Table S3. Removing moderation by SES of *ACE* influences on cortisol, cortisone, or cross-trait *ACE* correlation estimates each did not significantly decrease model fit relative to the baseline model, nor did removing all moderation by SES. Inspection of individual parameter estimates in the full moderation model (Table S5) revealed significant moderation of shared environmental effects by SES for both cortisol and cortisone (Figure S3). Nonparametric analyses indicated a trend of heightened genetic influence and reduced shared environmental influences on cortisol and cortisone at lower SES (Figure S4). This is consistent with a diathesis-stress hypothesis, which predicts that genetic influences are stronger under higher stress conditions. However, post-hoc multiple-group tests of these differences indicated that they were not statistically significant (Table S6). Even larger sample sizes than those implemented here may be necessary in order to test definitively for SES moderation.

## Discussion

In the largest twin study of neuroendocrine concentrations in any tissue type, we estimated genetic and environmental contributions to child and adolescent hair cortisol and cortisone, allowing for moderation by age, sex, and family socioeconomic status. We found moderate genetic influences on both hormones, with some indication of stronger shared environmental influences on male cortisone than on female cortisone. Genetic influences on cortisol decreased with advancing age, whereas genetic influences on cortisone increased with age.

This sample contained high levels of socioeconomic diversity, with one-third of families reporting having received means-tested public assistance since the twins' birth. Nevertheless, we did not find strong shared environmental influences on either cortisol or cortisone, nor did we find evidence that family SES moderates genetic and environmental influences on cortisol or cortisol, a commonly studied biomarker presumed to index chronic stress. Moreover, counter to expectations, we found no evidence for systematic associations between family socioeconomic status and mean levels of either cortisol or cortisone. It remains possible that the socioeconomic status affects HPA function by way of specific components of diurnal variation,^45^ without altering total cortisol output.

Previous large-scale studies of hair cortisol and cortisone have either focused on adults^25,41^ or on six year olds.^20^ The current study presents the first rigorous examination of age trends, sex differences, and SES differences in cortisol and cortisone over middle childhood and adolescence. We found marked sex differences in age trends. At age 8 years, females evince lower average levels of both hair cortisol and cortisone than males, but females increase in glucocorticoid concentrations over the course of adolescence more rapidly than males, such that by age 18 years females have slightly higher mean levels of both hormones. This finding may provide one plausible biological mechanism for the escalation of internalizing psychopathology in females over the course of adolescence.^46,47^

One previous large scale study reported a correlations of .55 between hair cortisol and cortisone in adults^41^, which is very similar to the values that we report in the current child and adolescent sample (.51 for males and .54 for females). Biometric decompositions indicated that approximately half of this correlation is attributable to shared genetic etiology. The source of the remaining, environmentally-driven covariation between cortisol-cortisone changed dynamically with age, shifting from family-level (shared) influences prior to ∼age 10 years, to twin-specific (non-shared) environments thereafter.

The opposing age trends in the magnitudes of genetic influence on cortisol and cortisone are noteworthy. One possibility is that, over the course of adolescent development, homeostatic processes increasingly dampen genetically-influenced variation in cortisol by converting it into inactive cortisone. These shifts also occur concomitantly with age-related decreases in shared environmental influences on both cortisol and cortisone, suggesting a more general buffering pattern of both genetic and shared environmental effects on cortisol with advancing age.

It is useful to consider possible mechanisms for genetic influence on HPA activity. Most obviously, polymorphisms in the genes involved in glucocorticoid synthesis, release, and metabolism may be related to homeostatic levels of circulating cortisol. However, although cortisol and cortisone are molecular phenotypes, the pathways between genotype and hormonal concentrations may be less direct. Genetic differences between people also shape the likelihood that they will experience stressful events. For example,neighborhood quality, life events, and relationship disruption have all been shown to be heritable, i.e., systematically associated with genetically-influenced individual differences.^48–50^ Additionally, genetic variation in other biological pathways, not directly involved in glucocorticoid metabolism, may shape psychological factors that, in turn, influence how stressful events are interpreted and coped with.Therefore, genetic influences on phenotypes measured “under the skin” may nevertheless be translated via environmental, “outside the skin” pathways.^51^ Finally, HPA axis changes may be an outcome of disease processes, such as major depression.^52^

In conclusion, this study is the first to estimate the magnitude of genetic influences on long-term glucocorticoid output and to examine how genetic influences differ with age, sex, and socioeconomic status. Further research spanning levels of measurement and explanation will be needed to understand the mechanisms of these genetic influences.

## Supplementary Information

Supplementary information is available online.

## Acknowledgements

This research was supported by National Institutes of Health (NIH) grants R01HD083613, R21HD081437, and R21AA023322. EMTD & KPH were supported as visiting scholars at the Russell Sage Foundation. LEE was supported by a National Science Foundation Graduate Research Fellowship. The Population Research Center at the University of Texas at Austin is supported by NIH grant R24HD042849.

### Conflict of Interest

The authors declare no conflict of interest.

## Supplementary Information

### Specification of Moderated Biometric Models

Models were specified as three-group (MZ, same-sex DZ, opposite-sex DZ) biometric decompositions of cortisol and cortisone into additive genetic (A), shared environmental (C), and non-shared environmental factors unique to each twin (E) in the form of a bivariate correlated factors model (Figure S1). Within a phenotype, the *A* factor is fixed to be correlated at 1.0 and 0.5 in MZ and same sex DZ twins, respectively. The *C* factor is, by definition, fixed to correlate at 1.0 in all same twin pairs. As *E* factor captures all variance not shared between MZ twins, including error variance, they are not correlated within phenotypes.

Qualitative sex differences are represented by allowing within-and across-phenotype A or C correlations to differ across same sex and opposite sex DZ twins. As these correlations cannot be estimated for additive genetic and shared environmental factors simultaneously,^1^ qualitative sex differences were examined for additive genetic and shared environmental factors separately. Models that do not allow for qualitative sex differences constrain within-and across-phenotype A and C correlations to be equal across same sex and opposite sex twin pairs. Quantitative sex differences are tested by allowing for moderation of *ACE* paths and *A* and/or *C* correlations by sex. Sex was effects coded (female = -.5, male = .5) such that the main effects parameters of A, C, and *E* represent population-mean effects (assuming an equal sex distribution in the population) and interaction effects parameters of sex by A, *C,* and *E* represent the sex difference in the parameter value. *ACE* × age and *ACE* × SES interactions were similarly estimated by allowing for moderation of *ACE* paths (act, cct, ect and acn, ccn, ecn) and cross phenotype *ACE* correlations (rA, rC, and rE) by either Age or SES, in the form p = po + p1*m, where p is the parameter and m is the moderator. All models included main effects of race (Caucasian coded as reference group) and sex-specific main effects of age. Age was centered at 8 years of age to reflect the lowest observed integer value in the sample.

The initial bivariate model allowed for quantitative, qualitative, and sex-specific mean age differences. Model fit was examined by sequentially removing: (i) qualitative sex differences, (ii) quantitative sex differences, and (iii) sex-specificity of differences. Nested models were compared using Satorra-Bentler scaled chi-square difference tests.^2^ Models were also compared using the Akaike Information Criterion (AIC)^3^ and the Bayesian Information Criterion (BIC).

To calculate unstandardized variance in each phenotype accounted for by its *ACE* factors (left and middle panels of Figures 2 and S3), the paths from the *ACE* factors factors to the phenotype (act, cct, and ect for cortisol and acn, ccn, ecn for cortisone) are squared. To calculate *ACE* contributions to phenotypic correlation (right panels of Figures 2 and S3), one calculates the products of the constituent paths in the respective the A-, C-, and E-mediated pathways.Thus, the *A* contribution to phenotypic correlation is act × ra × acn. Similarly, the *C* contribution is cct × rc × ccn, and the *E* contribution is ect × re × ecn.

To calculate proportions of variance in each phenotype accounted for by its *ACE* factors (Figure S2), the unstandardized variance accounted for is divided by the total variance. For cortisol this yields act^2^ / (act^2^ + cct^2^ + ect^2^), cct^2^ / (act^2^ + cct^2^ + ect^2^), and ect^2^ / (act^2^ + cct^2^ + ect^2^), for proportions of variances accounted for by *ACT*, *CCT*,and *ECT*, respectively. For cortisone, this yields acn^2^ / (acn^2^ + ccn^2^ + ecn^2^), ccn^2^ / (acn^2^ + ccn^2^ + ecn^2^), and ecn^2^ / (acn^2^ + ccn^2^ + ecn^2^), for proportions of variance accounted for by *ACN*, *CCN*, and *ECN*, respectively.

### Non-parametric Analyses: Local Structural Equation Modeling (LOSEM)

LOSEM is an analytic approach that provides locally weighted estimates of *ACE* parameters along a continuous moderator, such as age.^4^ LOSEM uses a weighting kernel and bandwidth that gives observations in closer proximity to the focal value of the moderator greater weight. LOSEM thereby provides a non-parametric estimate of moderated trends by estimating multiple structural equation models that change only with respect to the assigned focal value of the moderator. Age and SES were not included as parameters in the structural equation models, but rather as weighting variables.

***LOSEM results: Moderation by age***. To evaluate trends in *ACE* estimates for cortisol and cortisone across age, 101 models were estimated with the focal value of age set from 8 to 18 at intervals of 0.1. Results from this model are displayed in the top row of Figure S4. LOSEM findings were largely consistent with parametric estimates, with two noticeable differences. First, LOSEM results indicated that genetic effects on cortisone primarily increased from ages 8 to 13, followed by a leveling off and even slow decline. This is in contrast to parametric results that indicated genetic effects on cortisone increase consistently across age. Second, parametric results indicated that shared environmental effects on cortisone were moderate at age 8 and age 18, while minimal to non-existent at age 14. Conversely, LOSEM results indicated that shared environmental effects on cortisone were minor at age 8, followed by estimates near 0 across the remainder of the age range.

***LOSEM results: Moderation by SES***. Forty-one LOSEM models for SES were estimated with focal values set between −2 and 2 standard deviations at intervals of 0.1. Results from this model are displayed in the bottom row of Figure S4. Additive genetic influences were high at low levels of SES followed by a decrease through approximately 1 *SD* and a subsequent slight increase. Shared environmental influences on cortisone decreased through −1 *SD* followed by an increase until approximately 1 *SD* and then a decrease through 2 *SD*. Non-shared environmental influences on cortisone were consistent across SES. For cortisol, additive genetic effects decreased through −1 *SD* followed by a leveling off. Shared and non-shared environmental influences both increased around −1 *SD* followed by a leveling off.

### Results of Post-Hoc Analyses: Dichotomous Age and SES Moderation

***Moderation by age***. Inspection of age LOSEM results for cortisol and cortisone indicated one transition point at approximately 13 years of age. Two discrete groups were created with participants 13 years of age or younger coded as 0 and participants older than 13 coded as 1. The baseline model allowed for interactions between the dichotomized age or SES variable and *ACE* estimates for cortisol, cortisone and their association. All models also included mean sex and age differences and an age x sex interaction; age was entered as a continuous covariate. Model fit indices for dichotomous age moderation models are summarized in the top portion of Table S6. Including moderation by age of the association between shared environmental factors prohibited model convergence and this interaction term was excluded from all models. As with the continuous moderation results, removing age moderation of the remaining *ACE* parameters for cortisol, cortisone, or the cross-trait correlations significantly decreased model fit relative to the baseline model. Removing age moderation entirely also significantly decreased model fit relative to the baseline model.

***Moderation by SES***. Inspection of SES LOSEM results revealed a transition point at approximately –0.5 *SDs*. Two discrete groups were created for participants with SES scores at or below –0.5 *SDs* and participants above –0.5 SDs. The baseline model also included SES as a continuous covariate. Model fit indices for dichotomous SES moderation models are summarized in the bottom portion of Table S6. As with age, including moderation by SES of the association between shared environmental factors resulted in a series of models that failed to converge and, therefore, this interaction term was excluded for all analyses. Model comparisons revealed that removing SES moderation of remaining *ACE* parameters for cortisol, cortisone, or the cross-trait correlations did not significantly decrease model fit relative to the baseline mode. In addition, removing all moderation by SES did not significantly decrease model fit. Finally, allowing SES to moderate only shared environmental influences did not significantly increase model fit relative to a model that included no moderation.

---

**Table S1.**
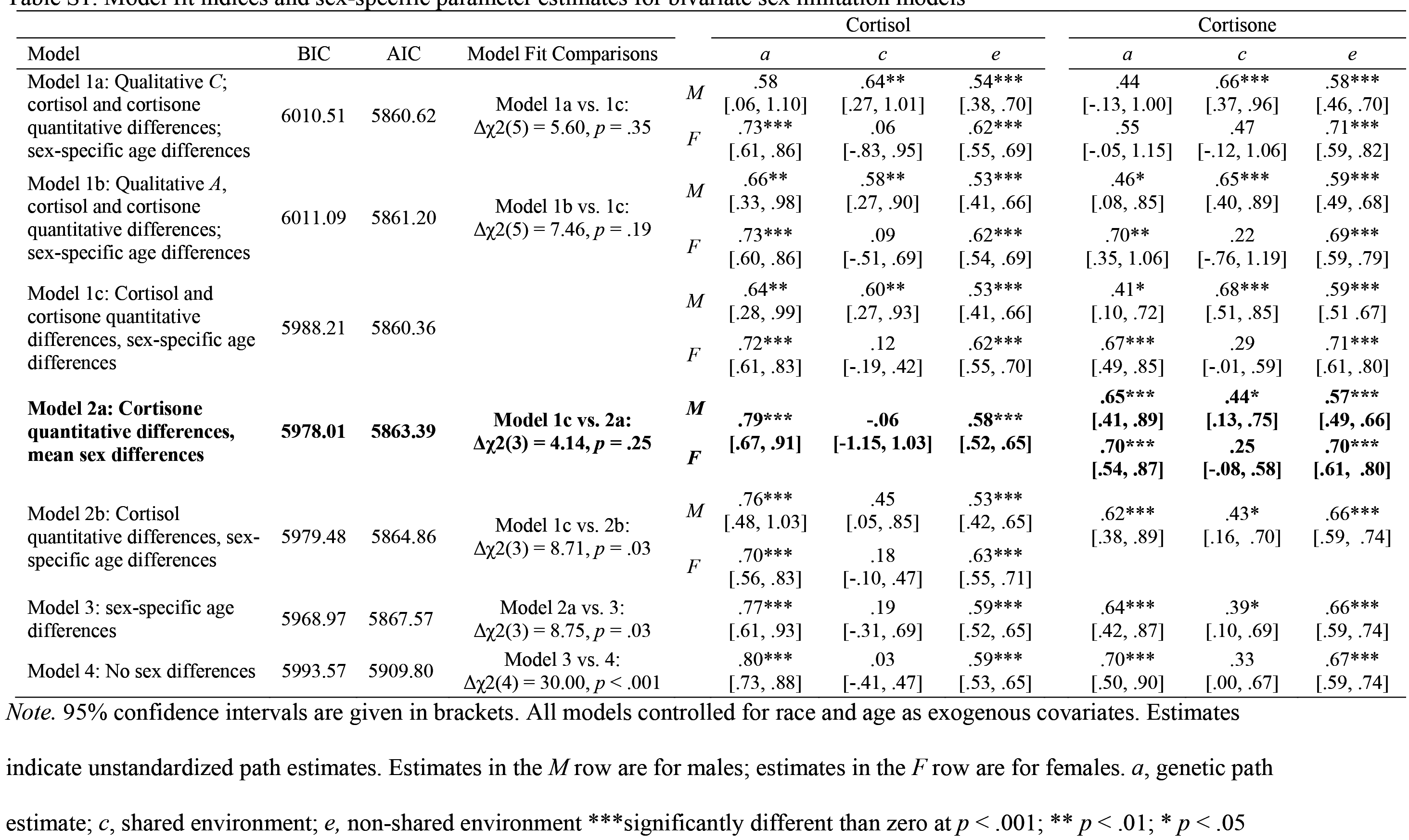
Model fit indices and sex-specific parameter estimates for bivariate sex limitation models

**Table S2.** Unstandardized Parameter Estimates from the Final Bivariate Sex LimitationModel

| Parameter | Estimate | (95% CI) | p-value |
| --- | --- | --- | --- |
| Variance in Cortisone |  |  |  |
| Main Genetic effect ($A_{CN}$) | .677 | (.491, .863) | < .001 |
| Gene × Sex Interaction ($A_{CN}'$) | -.054 | (-.245, .137) | .641 |
| Main Shared Environment effect ($C_{CN}$) | .346 | (.039, .652) | .064 |
| Shared Environment × Sex Interaction ($C_{CN}'$) | .190 | (.010, .370) | .083 |
| Main Non-shared Environmental effect ($E_{CN}$) | .638 | (.571, .705) | < .001 |
| Non-shared Environment × Sex Interaction ($E_{CN}'$) | -.130 | (-.250, -.010) | .074 |
| Main Sex effect | .631 | (.428, .833) | < .001 |
| Main Age effect | .082 | (.060, .104) | < .001 |
| Age × Sex Interaction | -.058 | (-.104, -.011) | .040 |
| Race (Hispanic) | .309 | (.126, .492) | .005 |
| Race (African American) | .334 | (.018, .649) | .082 |
| Race (Other) | .109 | (-.080, .299) | .344 |
| Variance in Cortisol |  |  |  |
| Main Genetic effect ($A_{CT}$) | .791 | (.669, .913) | < .001 |
| Main Shared Environment effect ($C_{CT}$) | -.059 | (-1.148, 1.030) | .929 |
| Main Non-shared Environment effect ($E_{CT}$) | .584 | (.523, .645) | < .001 |
| Main Sex effect | .568 | (.351, .785) | < .001 |
| Main Age effect | .049 | (.026, .072) | .001 |
| Age × Sex Interaction | -.065 | (-.106, -.024) | .009 |
| Race (Hispanic) | .020 | (-.130, .18-) | .827 |
| Race (African American) | .814 | (.442, 1.186) | < .001 |
| Race (Other) | -.083 | (-.292, .125) | .511 |

Note. 95% CI = 95% confidence interval. p-value = two-tailed probability of type-I error.

**Table S3.**
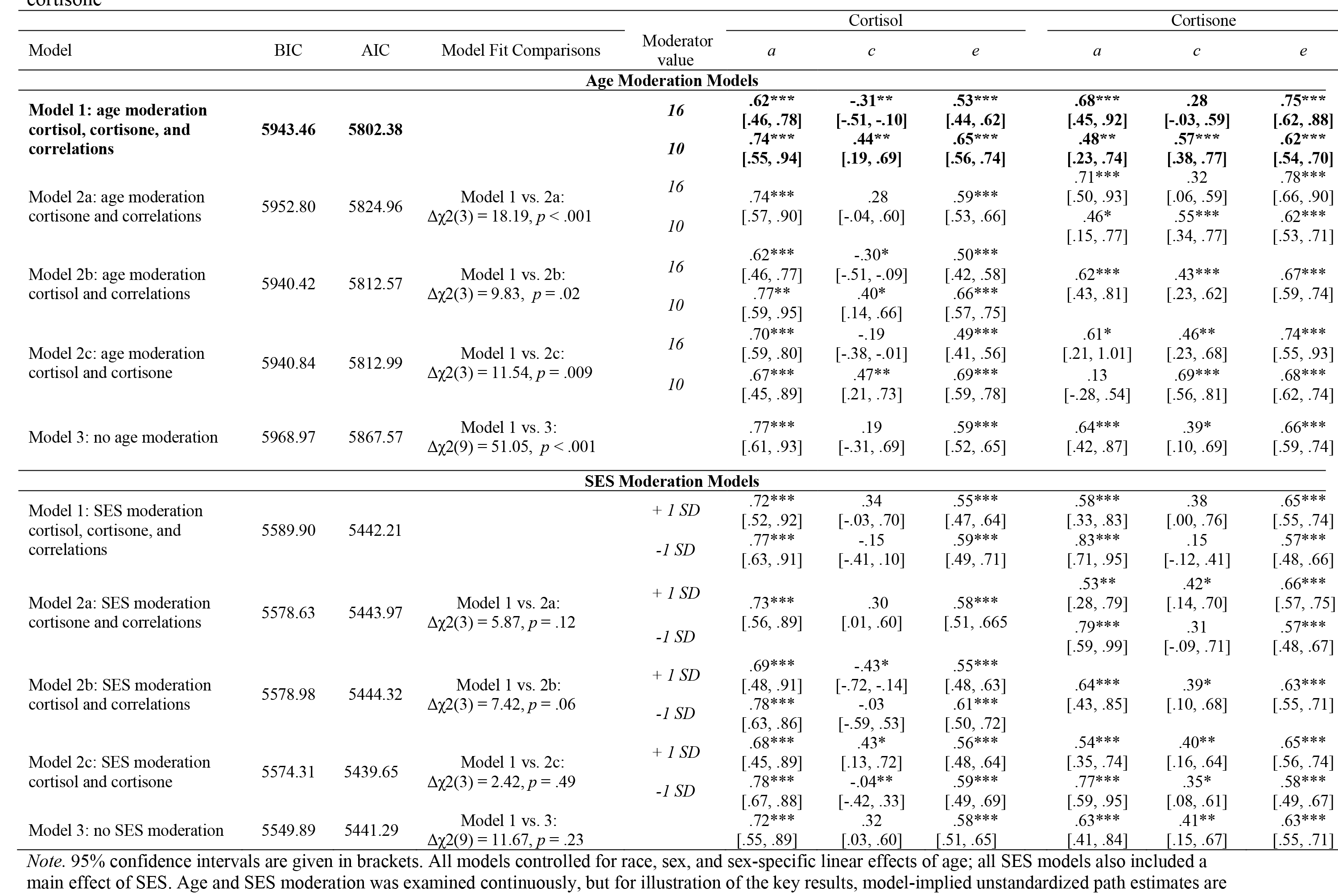
Model fits and simple slopes for bivariate age and SES moderation models of genetic and environmental influences on cortisol and cortisone

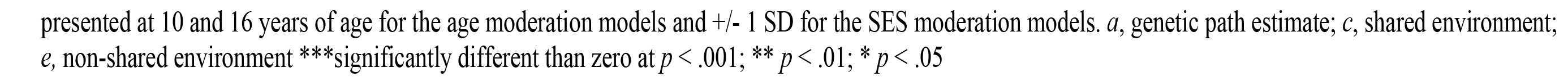

**Table S4.** Unstandardized Parameter Estimates from the Final Age Moderation Model

| **Parameter** | **Estimate** | **(95% CI)** | **p-value** |
| --- | --- | --- | --- |
| **Variance in Cortisone** |  |  |  |
| Main Genetic effect ($A_{CN}$) | .416 | (.090, .742) | .036 |
| Gene × Age Interaction ($A_{CN}'$) | .033 | (-.014, .081) | .251 |
| Main Shared Environment effect ($C_{CN}$) | .669 | (.418, .920) | < .001 |
| Shared Environment × Age Interaction ($C_{CN}'$) | -.049 | (-.102, .006) | .136 |
| Main Non-shared Environmental effect ($E_{CN}$) | .574 | (.461, .687) | < .001 |
| Non-shared Environment × Age Interaction ($E_{CN}'$) | .022 | (-.002, .046) | .128 |
| Main Sex effect | .618 | (.422, .813) | < .001 |
| Main Age effect | .082 | (.060, .104) | < .001 |
| Age × Sex Interaction | -.057 | (-.101, -.013) | .034 |
| Race (Hispanic) | .297 | (.111, .483) | .009 |
| Race (African American) | .332 | (.016, .648) | .084 |
| Race (Other) | .099 | (-.093, .290) | .397 |
| **Variance in Cortisol** |  |  |  |
| Main Genetic effect ($A_{CT}$) | .786 | (.499, 1.073) | < .001 |
| Gene × Age Interaction ($A_{CT}'$) | -.021 | (-.072, .030) | .491 |
| Main Shared Environment effect ($C_{CT}$) | .690 | (.382, .999) | < .001 |
| Shared Environment × Age Interaction ($C_{CT}'$) | -.125 | (-.165, -.047) | < .001 |
| Main Non-shared Environment effect ($E_{CT}$) | .687 | (.560, .815) | < .001 |
| Non-shared Environment × Age Interaction ($E_{CT}'$) | -.020 | (-.042, .003) | .156 |
| Main Sex effect | .539 | (.330, .748) | < .001 |
| Main Age effect | .051 | (.028, .074) | < .001 |
| Age × Sex Interaction | -.059 | (-.098, -.021) | .011 |
| Race (Hispanic) | .036 | (-.117, .190) | .697 |
| Race (African American) | .833 | (.468, 1.199) | < .001 |
| Race (Other) | -.077 | (-.266, .112) | .504 |
| **Cortisone with Cortisol** |  |  |  |
| Main Genetic correlation ($r_A$) | .735 | (.075, 1.396) | .067 |
| Gene × Age Interaction ($r_A'$) | .003 | (-.100, .106) | .960 |
| Main Shared Environment correlation ($r_C$) | .575 | (.089, 1.060) | .052 |
| Shared Environment × Age Interaction ($r_C'$) | -.099 | (-.273, .075) | .349 |
| Main Non-shared Environment correlation ($r_E$) | .186 | (-.025, .398) | .148 |
| Non-shared Environment × Age Interaction ($r_E'$) | .060 | (.030, .090) | .001 |

*Note.* 95% CI = 95% confidence interval. p-value = two-tailed probability of type-I error.

**Table S5.** Unstandardized Parameter Estimates from the Full SES Moderation Model

| Parameter | Estimate | (95% CI) | p-value |
| --- | --- | --- | --- |
| Variance in Cortisone |  |  |  |
| Main Genetic effect ($A_{CN}$) | .704 | (.547, .862) | < .001 |
| Gene × SES Interaction ($A_{CN}'$) | -.125 | (-.240, -.009) | .076 |
| Main Shared Environment effect ($C_{CN}$) | .263 | (-.054, .580) | .172 |
| Shared Environment × SES Interaction ($C_{CN}'$) | .115 | (.032, .199) | .023 |
| Main Non-shared Environmental effect ($E_{CN}$) | .608 | (.543, .673) | < .001 |
| Non-shared Environment × SES Interaction ($E_{CN}'$) | .036 | (-.030, .103) | .369 |
| Main SES effect | -.004 | (-.071, .063) | .920 |
| Main Sex effect | .564 | (.363, .765) | < .001 |
| Main Age effect | .075 | (.053, .097) | < .001 |
| Age × Sex Interaction | -.039 | (-.080, .001) | .110 |
| Race (Hispanic) | .212 | (.024, .399) | .064 |
| Race (African American) | .282 | (-.042, .606) | .153 |
| Race (Other) | .097 | (-.091, .284) | .396 |
| Variance in Cortisol |  |  |  |
| Main Genetic effect ($A_{CT}$) | .747 | (.650, .843) | < .001 |
| Gene × SES Interaction ($A_{CT}'$) | -.025 | (-.168, .117) | .770 |
| Main Shared Environment effect ($C_{CT}$) | .092 | (-.189, .373) | .592 |
| Shared Environment × SES Interaction ($C_{CT}'$) | .243 | (.103, .383) | .004 |
| Main Non-shared Environment effect ($E_{CT}$) | .574 | (.511, .636) | < .001 |
| Non-shared Environment × SES Interaction ($E_{CT}'$) | -.020 | (-.095, .055) | .655 |
| Main SES effect | .023 | (-.044, .089) | .578 |
| Main Sex effect | .542 | (.326, .759) | < .001 |
| Main Age effect | .048 | (.024, .072) | < .001 |
| Age × Sex Interaction | -.052 | (-.090, -.014) | .026 |
| Race (Hispanic) | .012 | (-.140, .165) | .895 |
| Race (African American) | .684 | (.284, 1.083) | .005 |
| Race (Other) | -.090 | (-.307, .127) | .496 |
| Cortisone with Cortisol |  |  |  |
| Main Genetic correlation ($r_A$) | .653 | (.437, .870) | < .001 |
| Gene × SES Interaction ($r_A'$) | -.029 | (-.383, .324) | .892 |
| Main Shared Environment correlation ($r_C$) | 1.329 | (.013, 2.644) | .097 |
| Shared Environment × SES Interaction ($r_C'$) | -.992 | (-2.170, .186) | .166 |
| Main Non-shared Environment correlation ($r_E$) | .407 | (.287, .528) | < .001 |
| Non-shared Environment × SES Interaction ($r_E'$) | -.002 | (-.166, .162) | .986 |

Note. 95% CI = 95% confidence interval. p-value = two-tailed probability of type-I error.

**Table S6.** Dichotomous Age and SES Moderation Models

| Model | BIC | AIC | Model Fit Comparisons |
| --- | --- | --- | --- |
| **Age Moderation** |  |  |  |
| **Model 1: age moderation cortisol, cortisone, and genetic and non-shared environmental correlations** | **5928.67** | **5792.00** |  |
| Model 2a: age moderation cortisone and correlations | 5952.80 | 5824.96 | Model 1 vs. 2a:<br>$\Delta\chi^2(3) = 16.23, p = .001$ |
| Model 2b: age moderation cortisol and correlations | 5925.95 | 5802.51 | Model 1 vs. 2b:<br>$\Delta\chi^2(3) = 10.33, p = .02$ |
| Model 2c: age moderation cortisol and cortisone | 5927.95 | 5800.11 | Model 1 vs. 2c:<br>$\Delta\chi^2(2) = 8.21, p = .02$ |
| Model 3: no age moderation | 5968.97 | 5867.57 | Model 1 vs. 3:<br>$\Delta\chi^2(8) = 61.12, p < .001$ |
| **SES Moderation** |  |  |  |
| Model 1: SES moderation cortisol, cortisone, and correlations | 5592.40 | 5449.05 |  |
| Model 2a: SES moderation cortisone and correlations | 5579.32 | 5449.01 | Model 1 vs. 2a:<br>$\Delta\chi^2(3) = 4.41, p = .22$ |
| Model 2b: SES moderation cortisol and correlations | 5576.01 | 5446.69 | Model 1 vs. 2b:<br>$\Delta\chi^2(3) = 1.97, p = .58$ |
| Model 2c: SES moderation cortisol and cortisone | 5581.12 | 5446.46 | Model 1 vs. 2c:<br>$\Delta\chi^2(2) = 1.10, p = .58$ |
| **Model 3: no SES moderation** | **5549.89** | **5441.29** | **Model 1 vs. 3:**<br>**$\Delta\chi^2(8) = 5.17, p = .74$** |
| Model 4: SES moderation shared environment | 5562.09 | 5444.80 | Model 1 vs. 4:<br>$\Delta\chi^2(6) = 4.96, p = .55$<br>Model 3 vs. 4:<br>$\Delta\chi^2(2) = 0.28, p = .87$ |

*Note.* All models controlled for race, sex, and sex-specific linear effects of age; all SES models also included a main effect of SES.

**Figure S1.**
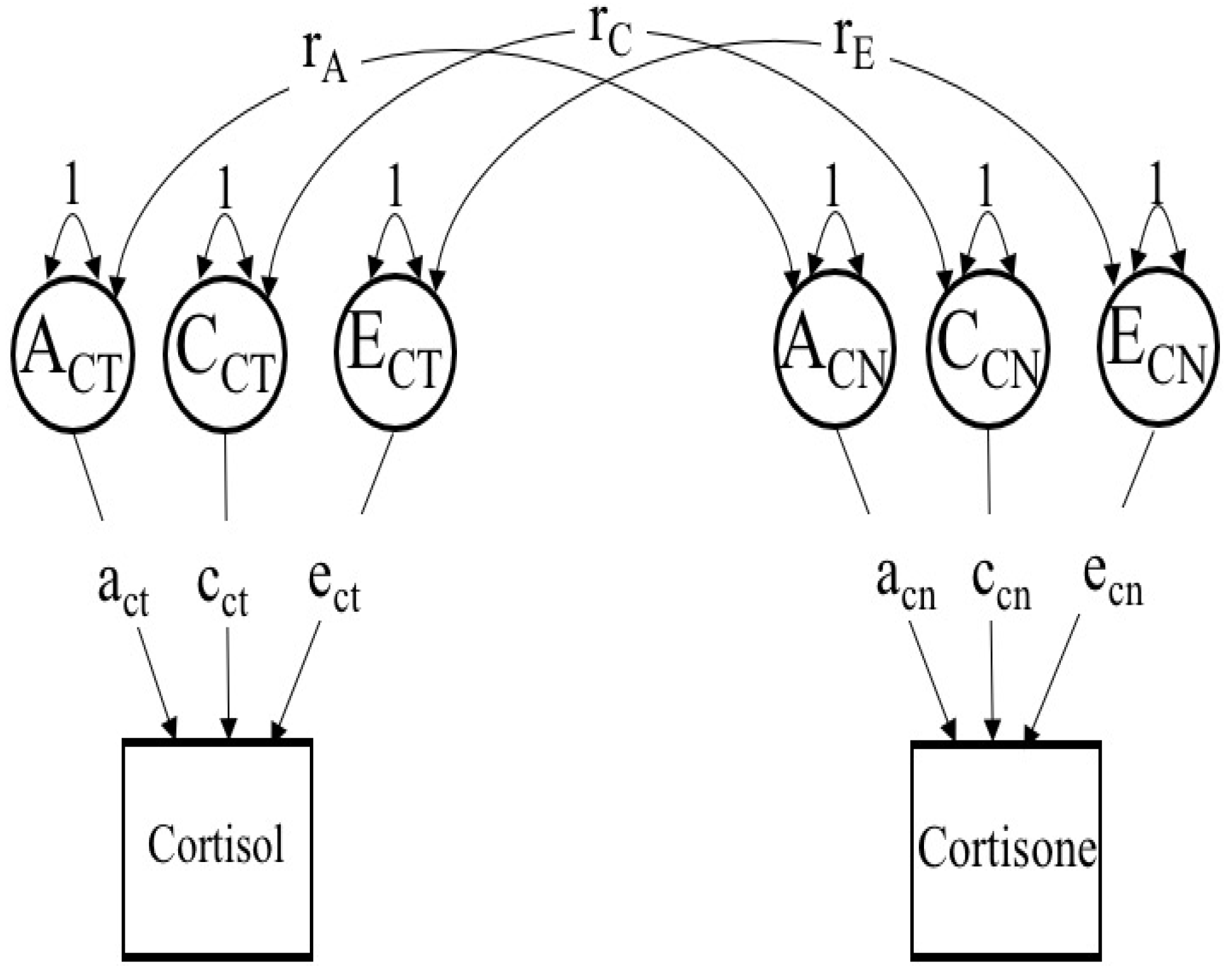
Path diagram for bivariate correlated factors model used for all parametric, biometric analyses. For ease of presentation only one twin per pair is depicted.

**Figure S2.**
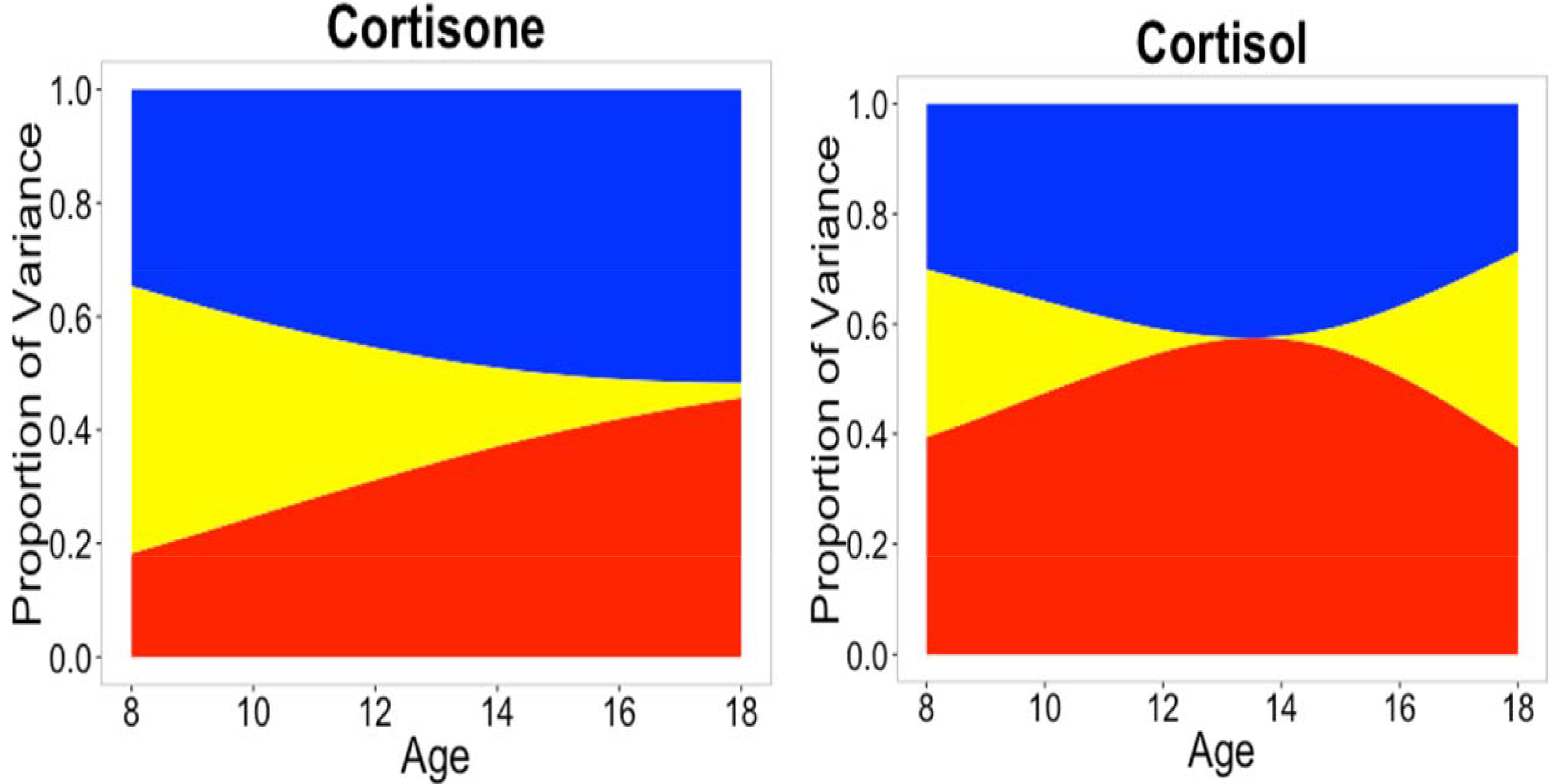
Proportions of variance explained by the ACE components in cortisone and cortisol as moderated by age.

**Figure S3.**
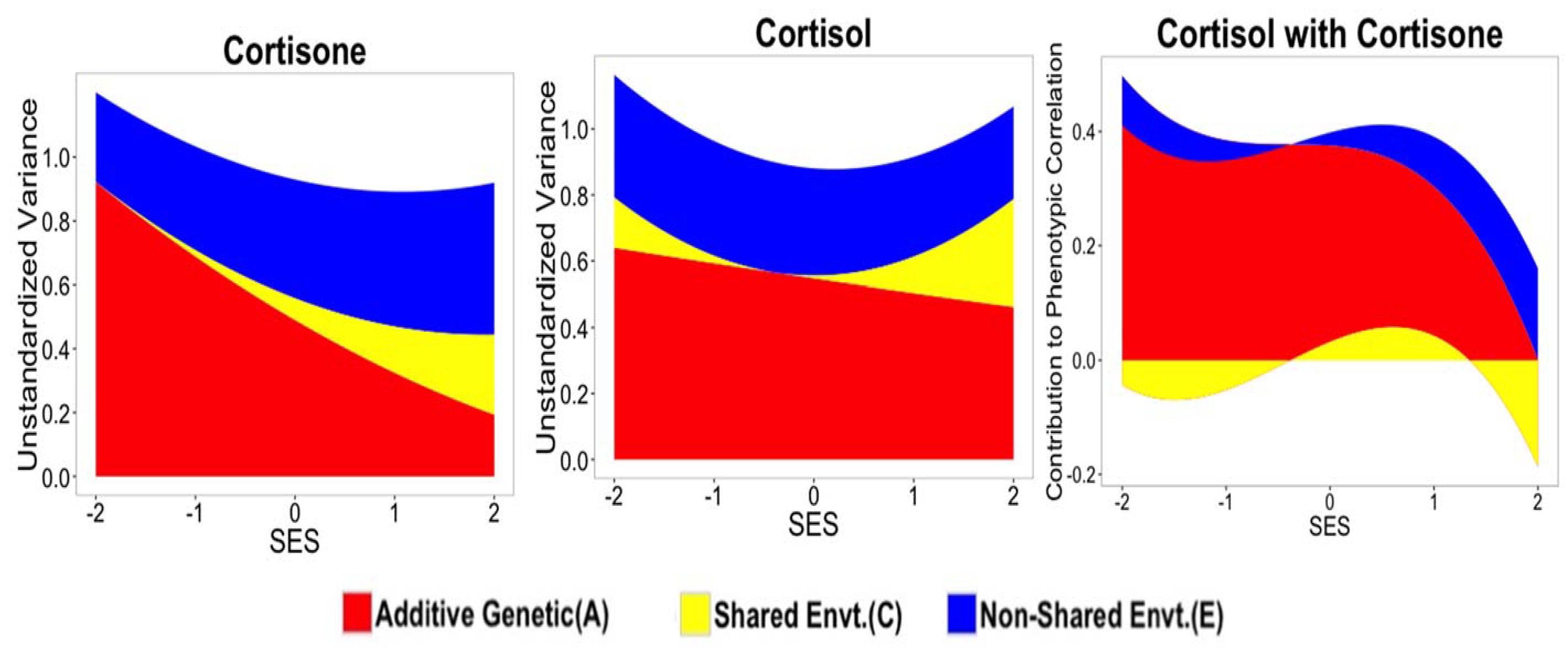
Moderation by SES of *ACE* contributions to unstandardized variance in cortisone and cortisol and their bivariate association.

**Figure S4.**
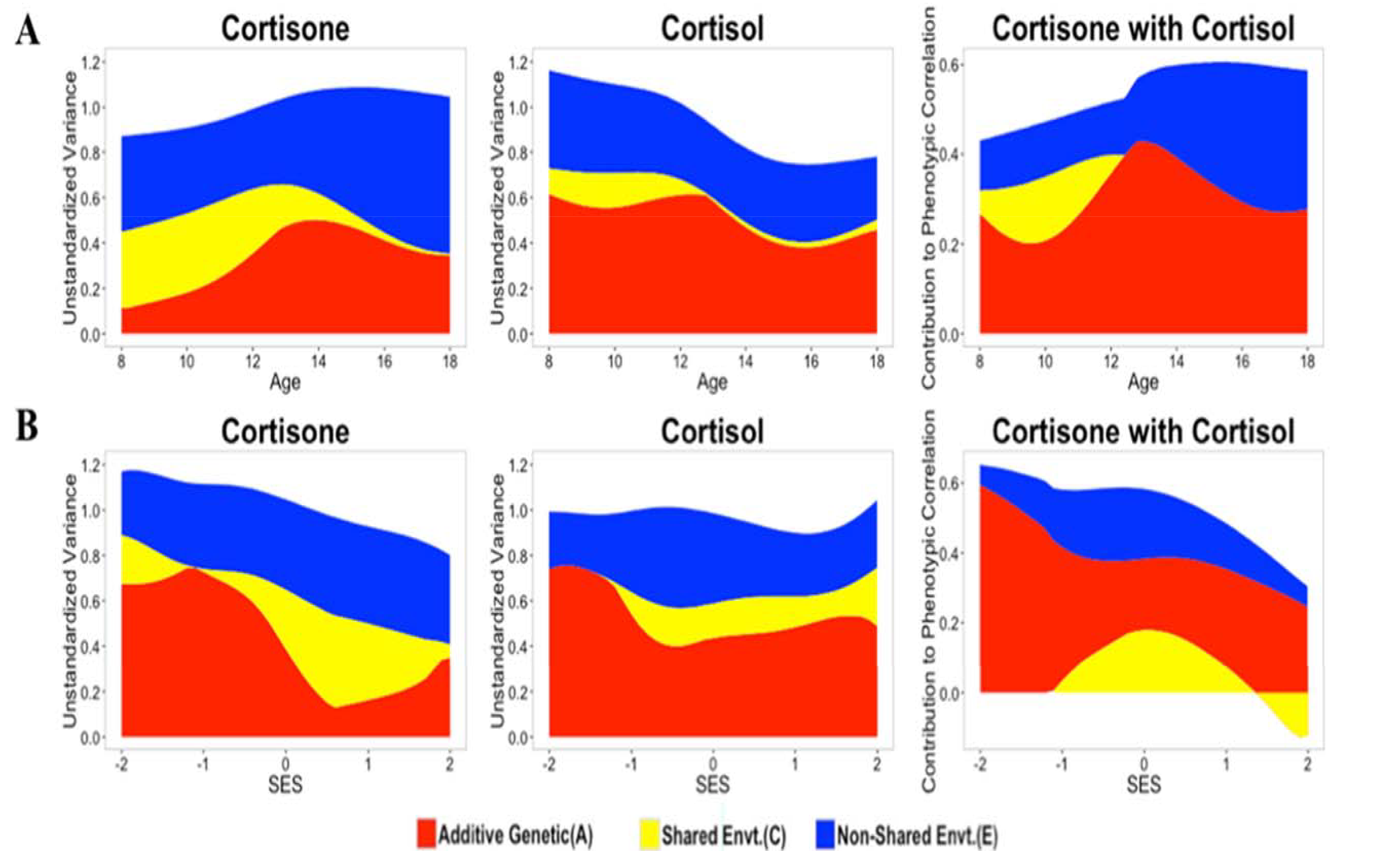
Unstandardized *ACE* estimates for cortisol and cortisone and *ACE* contributions to their bivariate association across age (row A) and SES (row B) using LOSEM.Shared environmental contributions to phenotypic correlations are not depicted when shared environmental variance was estimated at < .01 for cortisol or cortisone.

